# The human canonical core histone catalogue

**DOI:** 10.1101/720235

**Authors:** David Miguel Susano Pinto, Andrew Flaus

## Abstract

Core histone proteins H2A, H2B, H3, and H4 are encoded by a large family of genes distributed across the human genome. Canonical core histones contribute the majority of proteins to bulk chromatin packaging, and are encoded in 4 clusters by 65 coding genes comprising 17 for H2A, 18 for H2B, 15 for H3, and 15 for H4, along with at least 17 total pseudogenes. The canonical core histone genes display coding variation that gives rise to 11 H2A, 15 H2B, 4 H3, and 2 H4 unique protein isoforms. Although histone proteins are highly conserved overall, these isoforms represent a surprising and seldom recognised variation with amino acid identity as low as 77% between canonical histone proteins of the same type. The gene sequence and protein isoform diversity also exceeds commonly used subtype designations such as H2A.1 and H3.1, and exists in parallel with the well-known specialisation of variant histone proteins. RNA sequencing of histone transcripts shows evidence for differential expression of histone genes but the functional significance of this variation has not yet been investigated. To assist understanding of the implications of histone gene and protein diversity we have catalogued the entire human canonical core histone gene and protein complement. In order to organise this information in a robust, accessible, and accurate form, we applied software build automation tools to dynamically generate the canonical core histone repertoire based on current genome annotations and then to organise the information into a manuscript format. Automatically generated values are shown with a light grey background. Alongside recognition of the encoded protein diversity, this has led to multiple corrections to human histone annotations, reflecting the flux of the human genome as it is updated and enriched in reference databases. This dynamic manuscript approach is inspired by the aims of reproducible research and can be readily adapted to other gene families.

## Introduction

Histones are among the most abundant proteins in eukaryotic cells and contribute up to half the mass of chromatin [1]. The core histone types H2A, H2B, H3, and H4 define the structure and accessibility of the nucleosome as the fundamental repeating unit of genome organisation around which the DNA is wrapped [44]. In addition, the many chemically reactive sidechains of histones are post-translationally modified as a nexus for signalling and heritable epigenetics [40].

Core histones are delineated as either canonical or variant based on their gene location, expression characteristics, and functional roles (Table 1). Canonical core histones contribute the majority of proteins to the bulk structure and generic function of chromatin, and are encoded by 82 genes in 4 clusters named HIST1-HIST4 in the human genome, of which 65 are coding genes and 17 are pseudogenes (Table 2).

**Table 1:** Properties distinguishing canonical and variant core histone proteins.

|  | Canonical | Variants |
| --- | --- | --- |
| Expression timing | Replication dependent | Replication independent |
| Sequence identity | High | Low |
| Functional relationships | Isoforms | Specialised functions |
| Transcript stabilisation | Stem-loop | poly(A) tail |
| Gene distribution | Clusters | Scattered |

**Table 2:** Count of human canonical core histone coding genes and pseudogenes by histone cluster and type. ψ indicates pseudogenes.

|  | H2A | H2B | H3 | H4 | Total |
| --- | --- | --- | --- | --- | --- |
| HIST1 | 12 + 6ψ | 15 + 3ψ | 10 + 1ψ | 12 + 1ψ | 49 + 11ψ |
| HIST2 | 4 + 0ψ | 2 + 4ψ | 4 + 1ψ | 2 + 0ψ | 12 + 5ψ |
| HIST3 | 1 + 0ψ | 1 + 1ψ | 1 + 0ψ | 0 + 0ψ | 3 + 1ψ |
| HIST4 | 0 + 0ψ | 0 + 0ψ | 0 + 0ψ | 1 + 0ψ | 1 + 0ψ |
| Total | 17 + 6ψ | 18 + 8ψ | 15 + 2ψ | 15 + 1ψ | 65 + 17ψ |

Relationships within the histone family have been described using a variety of terminologies reflecting biochemical, functional, and genomic perspectives that are briefly described below and summarised in Table 3.

**Table 3:**
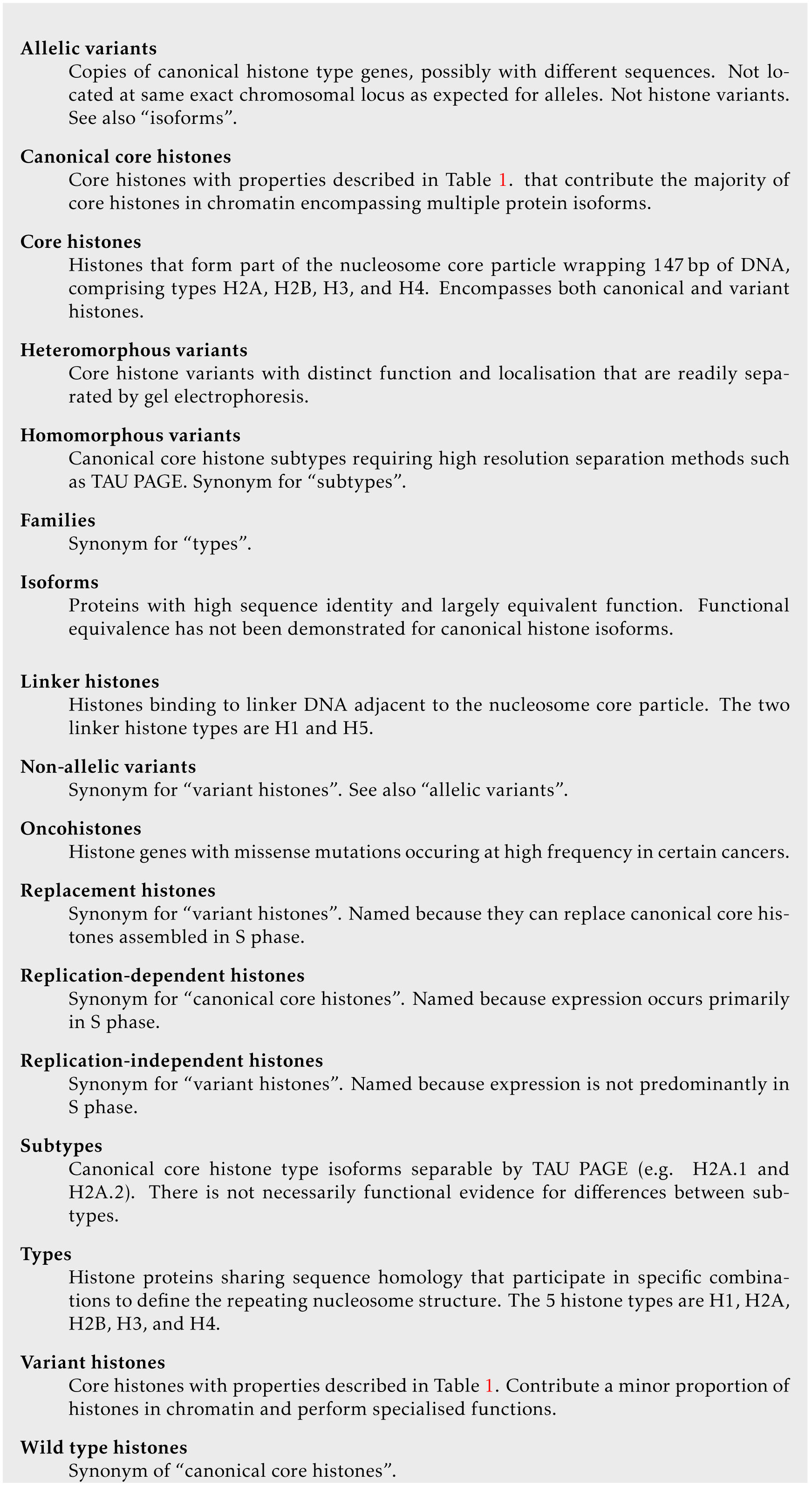
Terminology describing histone variation.

### Biochemical perspective

Abundant histone proteins are readily isolated using their highly basic chemical character. Successive improvements in fractionation ultimately revealed 5 main histone types with nomenclature H1, H2A, H2B, H3, and H4 [9]. An additional H1-related histone H5 is recognised in avian erythrocytes [41].

The demonstration of the nucleosome as the fundamental repeating unit of chromatin [39] showed that H2A, H2B, H3, and H4 associate as an octamer of two copies each within the nucleosome core particle. These four histones are referred to as core histones. In contrast, H1 associates with the linker DNA between nucleosome core particles and is referred to as a linker histone. The somatic H1 isoforms and tissue-specific variants are described elsewhere [31].

Arginine and lysine content was used as an early distinction between the histones [18]. The H1 linker histone has a low arginine/lysine ratio of 0.10 and became known as lysine-rich whereas the 4 core histones are arginine-rich with high arginine/lysine ratios of 0.91 in H2A, 0.40 in H2B, 1.38 in H3, and 1.25 in H4 type isoforms. Nevertheless, the core histones contain many lysines particularly in their N-terminal tails.

Separating histones by polyacrylamide gel electrophoresis (PAGE) using the strongly anionic detergent sodium dodecyl sulphate and neutral buffers (SDS PAGE) gives single bands for each histone type [66]. However, PAGE with non-ionic detergent Triton X–100 and urea as denaturants in acid buffers (TAU or AUT PAGE) allows the separation of histone types into multiple bands due to post-translational modifications and differences at specific amino acids in the polypeptides [87]. These TAU PAGE separations gave rise to subtype designations H2A.1, H2A.2, H3.1, H3.2, and H3.3.

### Functional perspective

Canonical core histone expression is significantly elevated during S phase to provide chromatin packaging for DNA duplicated during replication [81]. This led to their description as “replication dependent”, although a supply of canonical histones is inevitably required to partner variants throughout the cell cycle. Metazoan canonical core histone genes are distinctive because they lack introns and give rise to non-polyadenylated protein coding transcripts. Transcript turnover is regulated via a highly conserved 3’ stem-loop [50] (Table 1).

In contrast, variant histones such as H2A.Z, TH2B, H3.3, and CENP-A have reduced sequence identity and lower abundance [76]. They play functionally specific roles and are mostly expressed outside S phase, so are described as “replication independent”. Since histone variants are interpreted as taking the place of equivalent canonical core histone types, they are referred to as “replacement” histones.

### Genomic perspective

Canonical core histone genes are found in 4 clusters. The multiple gene copies in these clustered arrays are sometimes confusingly referred to as “allelic” and the resulting combined protein isoforms are often considered to be “wild type” although both genes and protein products display variation in primary sequence and relative abundance, and their functional equivalence has not been tested. A number of mutations in individual canonical and variant histone genes have been revealed to contribute to cancer [12, 21]. These “oncohistones” demonstrate that single amino acid changes in individual histone genes can have functional implications. In contrast, almost all variant histones are encoded by single genes dispersed in the genome with typical properties including introns, alternative splicing, and polyadenylated transcripts (Table 1).

### Cataloguing of canonical core histone diversity

Despite the importance of histones for chromatin organisation and extensive interest in their role in epigenetics and regulation, the curation and classification of human histone gene and protein sequences has not been systematically revisited since the landmark survey by *Marzluff et al*. [49]. The Histone Database has for many years provided an online database of histone sequences from across eukarya using sequence homology searching [5, 6, 7, 45, 73, 71, 72, 47, 48, 15], while some manually collated listings including human histones have also been undertaken [16, 37, 36]. This reflects continual revisions in genome sequence databases and the ongoing need for information about human histones. In fact, differences in 38 canonical core histone gene details (Table S1) have so far accumulated since the original 2002 survey due to rich annotations continuing to propagate into reference sequence databases.

In this manuscript we provide a comprehensive catalogue of canonical core histone genes, encoded proteins, and pseudogenes using reference genome annotations as the originating source. This reveals a surprising and seldom recognised variation in encoded histone proteins that exceeds commonly used subtype designations but whose functional implications have not been investigated.

Since curation and annotation of reference databases are dynamic and evolving, we have implemented the manuscript so that it can be regenerated at any time from the most current data in the NCBI RefSeq database in order to maintain its ongoing value as a reference source in an accessible format. All figures and tables were automatically generated using NCBI RefSeq data from 25th July 2019. All dynamically generated values in this copy of the text are displayed with a light grey background. The manuscript generation process has remained stable in our laboratory for several years and represents an example of “reproducible research” [64] that provides a novel model for cataloguing gene families.

## Histone genes

### Canonical core histone gene nomenclature

Canonical core histone genes adhere to a Human Genome Organisation (HUGO) Gene Nomenclature Committee (HGNC) endorsed system derived from the cluster number and position relative to other histones [49]. This superseded an earlier arbitrary scheme with backslashes (e.g. H2b/b) that preceded genome sequencing [3, 2].

The canonical core histone gene symbols are divided into 3 parts: HIST cluster, histone type, and identifier letter for the order relative to other histone genes of the same type in the same cluster (Figure 1(a)). For example, *HIST1H2BD* is nominally the fourth H2B coding gene in the HIST1 cluster. Identifiers are ordered by their genomic coordinates starting at the telomere of the short arm [49].

**Figure 1:**
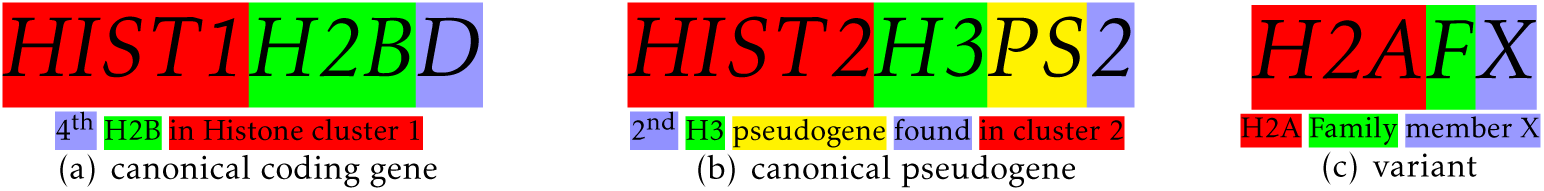
Histone gene nomenclature. (a) Canonical core histone gene names encode relative genomic order by cluster. (b) Pseudogenes named since 2002 include cluster, PS label, and discovery order identifier. (c) Most variant core histone genes are identified by type then F for family and identifier letter.

Two exceptions were originally applied to these simple naming rules [49]. Firstly, the positional identifier is omitted if there are no other histones of the same type in the cluster, so *HIST3H2A* is the sole H2A gene in HIST3. Secondly, the human and mouse histone clusters are largely syntenic so positional identifier letters for missing orthologs were skipped to maintain the equivalence of gene symbols. Consequently there is no human *HIST1H2AF* to accommodate *Hist1h2af* in mouse while keeping both –E and –G identifiers consistent for mouse and human orthologs.

Several new histone genes have been uncovered since the original naming (Table S1) and this required additional nomenclature exceptions. For example, new H2A encoding genes were identified in human and mouse HIST2 clusters preceding *HIST2H2AA*/*Hist2h2aa*. This led to the renaming of the gene to *HIST2H2AA3*/*Hist2h2aa3* and the addition of a new human gene as *HIST2H2AA4*. There are no human orthologs of the new mouse *Hist2h2aa1* and *Hist2h2aa2*.

Furthermore, no distinction was originally made between pseudogenes and functional coding genes, so *HIST3H2BA* is a pseudogene whereas neighbouring *HIST3H2BB* is the only functional H2B coding gene in HIST3. The HGNC definition of a pseudogene is a sequence that is generally untranscribed and untranslated but which has at least 50% predicted amino acid identity across 50% of the open reading frame to a named gene [26]. Newly uncovered histone pseudogenes are now suffixed with PS and a number in order of discovery (Figure 1(b)), such as *HIST1H2APS5* as the fifth H2A pseudogene discovered in HIST1. However, the pseudogenes named in the original classification retain their symbols without PS. This means that the absence of a PS suffix does not indicate a functional gene, and that there is no positional information in the gene symbols of most pseudogenes. Recently, some pseudogenes have also been shown to be functional histone variants [74].

### Histone gene clustering

The human canonical core histone genes are located in clusters HIST1 to HIST4, named in order of decreasing histone gene count (Table 2). HIST1 is the major histone gene cluster at locus 6p22.1–6p22.2 with 49 functional core histone genes plus all canonical H1 histones, representing 75% of all canonical core histone genes. HIST2 at locus 1q21.1–1q21.2 contains 12 coding genes, HIST3 at locus 1q42.13 contains 3 coding genes, and HIST4 at locus 12p12.3 contains 1 coding gene.

HIST1 and HIST2 are both contiguous high density arrays of histone genes. HIST1 spans 15.0 Mbp and is the second most gene dense region in the human genome at megabase scale after the MHC class III region [83]. The only non-histone protein coding gene located within the principal region of this cluster is *HFE*, encoding the hemochromatosis protein [4], although a number of other genes are located proximal to the outlying *HIST1H2AA* and *HIST1H2BA* pair.

It has also been argued that histone clustering does not contribute to gene conversion [54]. HIST1 is located towards the distal end of the major histocompatibility complex (MHC) in the extended class I region [28, 78] and it has even been suggested that this proximity may suppress recombination due to the observed local linkage disequilibrium [29]. In contrast, HIST2 may be prone to deletions and frequent rearrangements [65, 10]

The functional significance of histone gene clustering remains to be demonstrated. It has been suggested that clustering may facilitate coordinate regulation [17, 57], so interpreting such a functional relationship with genome organisation requires an accurate catalogue of the histone genes and their individual roles.

Conversely, progress in understanding histone gene function suggests there is a need to update the definitions of the canonical histone gene clusters. For example, the *HIST1H2AA* and *HIST1H2BA* genes are located 300 kbp upstream of the rest of the HIST1 cluster and separated from other cluster members a number of non-histone genes. Although the two genes have been assigned canonical histone gene names, the *HIST1H2BA* gene in fact encodes the most divergent canonical H2B protein which is also known as TH2/TH2B/hTSH2B and considered to be a testes-specific histone protein [86, 43, 67]. Immediately adjacent is *HIST1H2AA* encoding a H2A protein isoform of similarly high variation. The syntenic rat orthologues of *HIST1H2AA* and *HIST1H2BA* are divergently transcribed specifically in testes [34], and the mouse orthologues H2AL1/TH2a and TH2B have been shown to participate in gametogenesis [25] as well as to enhance stem cell reprogramming [68, 59]. It is therefore possible that *HIST1H2AA* and *HIST1H2BA* are undergoing sub-functionalisation and could be reclassified as variant genes encoding histone proteins H2A.L and H2B.1 [75]. This would in turn require gene renaming and a recalculation of the length of the HIST1 cluster.

A similar case of updates in definition may apply for the small HIST3 cluster 80 Mbp downstream of HIST2. HIST3 contains protein coding genes *HIST3H2A, HIST3H2BB*, and *HIST3H3. HIST3H3* encodes testes-specific H3T/H3.4 that has distinctive biochemical properties and appears to be a histone variant [80, 42], while the proteins encoded by *HIST3H2A* and *HIST3H2BB* are amongst the most divergent canonical histones of their types.

The examples of the *HIST1H2AA* and *HIST1H2BA* pair and the HIST3 cluster illustrate the evolving nature of the human canonical histone complement. This demonstrates the need for a dynamic approach to classification and presentation.

### Histone gene sequences

Despite the high conservation of canonical core histone protein sequences, the coding regions of these genes exhibit considerable variation (Table S3, S4, S5, and S6).

These differences are largely located in the third base position of codons. The mean number of synonymous substitutions per site (*d*_*S*_) in the sets of histone gene type isoforms is 3.5 for H2A, 1.7 for H2B, 2.6 for H3, and 2.8 for H4 (Table S8). This is consistent with observations that synonymous codon divergence far exceeds non-synonymous variation for histone genes across eukaryotes [61, 63]. It supports a hypothesis that histone protein sequence conservation results from birth and death evolution through strong selective pressure at the protein level [54]. Despite the level of synonymous substitution, codon usage is strongly biased towards the most frequently used human codons for most amino acids (Table S9 and data not shown). This suggests that histone translation may be sensitive to tRNA abundance.

### Histone variant genes

The 32 annotated human histone variant genes are listed in Table S10 for completeness. Histone variant gene families comprise only one or a few copies dispersed across the genome outside the canonical histone clusters. For example, the three H3.3 variant encoding genes are located on chromosomes 1, 12, and 17, far removed from other histone genes.

Most variant gene symbols have a separate 3 part nomenclature consisting of histone type, F for family, and an identifier letter (Figure 1(c)). Increasing interest in histone variant function [51] coupled with a variety of usages and conflicts between species has led to guidelines for improved consistency in histone variant nomenclature [75].

## Histone transcripts

Canonical core histone transcripts are among the few protein coding messages that are not polyadenylated. Instead they rely on a unique 3’ UTR stem-loop structure for regulation of transscript stability. They do have a 5’ 7–methyl–guanosine cap [50].

Canonical core histone gene transcription is regulated by cell-cycle dependent phosphorylation of the histone-specific Nuclear Protein of the Ataxia-Telangiectasia locus (NPAT) coactivator and interaction with the accessory protein FADD-Like interleukin-1β-converting enzyme/caspase-8-ASsociated Huge protein (FLASH), resulting in assembly of histone locus bodies coordinating factors responsible for transcription and processing [50, 62, 32]. Variability is observed in canonical core histone isoform gene transcription, both by analysis of non-polyadenylated transcripts [85] and RNA polymerase II promoter occupancy [16].

Overall there is estimated to be a 35 fold increase in mammalian canonical histone transcripts during S phase, principally as a result of a 10 fold increase in mRNA stabilisation via stem-loop dependent mechanisms, and a 3–5 fold up-regulation in canonical core histone gene transcription [30].

This post-transcriptional regulation is achieved by a stem-loop encoded in the mRNA 3’ untranslated region that is recognised by spliceosome-related RNA processing and stabilisation complexes [77]. The start location of the annotated stem-loops in human canonical histone transcripts ranges from 22 to 67 bp after the stop codon with a modal value of 35 bp. The sequence logo of aligned stem-loops confirms that the stem-loop is highly conserved (Figure 2(a)).

**Figure 2:**
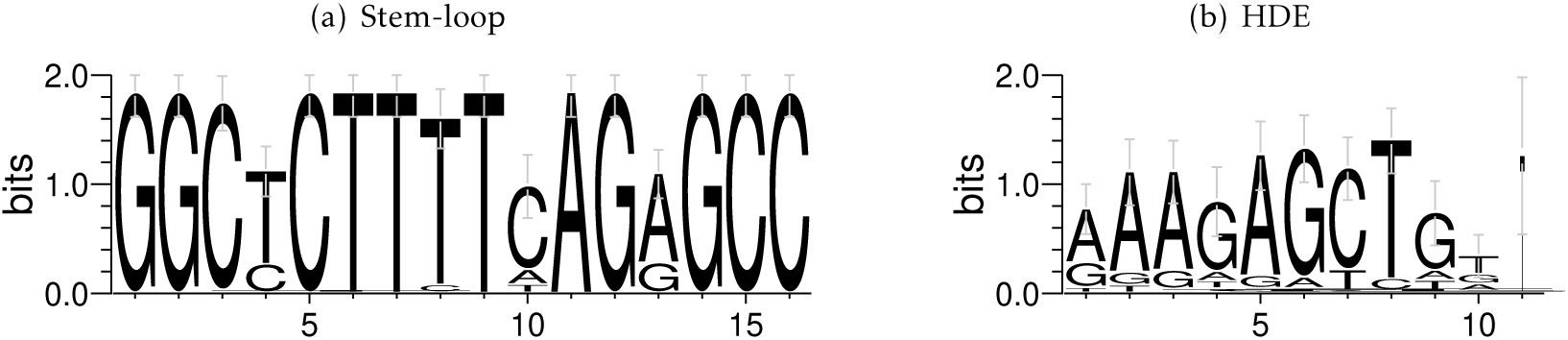
Sequence logos for (a) annotated stem-loops and (b) Histone Downstream Elements (HDEs) identified by homology for all canonical core histone gene 3’ untranslated regions (UTRs).

The RNA stem-loop structure is bound by the Stem-Loop Binding Protein (SLBP) which is up-regulated 10–20 fold during S phase to stabilise histone mRNAs [79]. Immediately downstream of the stem-loop a purine-rich Histone Downstream Element (HDE) interacts with U7 snRNA to direct efficient 3’ end processing. Although this is not an annotated feature of histone genes, alignment of the canonical HDE [23] to all canonical histone genes shows the modal location of the HDE is 16 bp downstream of the stem-loop. The sequence logo confirms that this feature is also highly conserved (Figure 2(b)).

Stabilisation and processing extend the half life of canonical core histone mRNAs during S phase and contribute to increased histone translation efficiency, enabling rapid production of histones to package the newly duplicated genomes.

Although the vast bulk of canonical core histone transcripts appear to be regulated by this mechanism, 1-5% of transcripts are found to be 3’ polyadenylated [85] and some genes have annotations indicating two transcripts differing in whether they have stem-loop or polyadenylation signals (Table S11).

Core histone variant transcripts lack a 3’ stem-loop and are polyadenylated in the same way as most protein coding genes. The exception is H2AX, which has alternatively processed transcripts exhibiting both the stem-loop characteristic of a canonical core histone and a poly(A) tail found on variant core histones [46, 60].

## Histone proteins

Many depictions of chromatin imply that canonical core histone protein types behave as a single “wild type” protein. This common assumption is based on the relatively high identity of histone sequences between isoforms and a focus of interest in functional roles of histone variants.

However, the very strong selection pressure on amino acid sequences encoded by canonical core histone genes [54], the roles of specific amino acid differences in observed proteins [51], and the consequences of small variations in proteins on the structure of nucleosomes [42], all suggest that the minor differences in the encoded canonical core histone protein isoforms could have functional implications.

Encoded isoform variation in canonical histone types H2A and H2B is pronounced. 11 H2A and 15 H2B distinct polypeptides are encoded by 17 H2A and 18 H2B coding genes, repectively. In contrast, only 4 H3 and 2 H4 distinct polypeptides are encoded by 15 H3 and 15 H24 coding genes repectively (Table 2). Although most variation is a result of amino acid substitutions, some H2A and H2B isoforms also show length differences.

The nomenclature for histone protein isoforms follows directly from the gene names [49], and supersedes the earlier nomenclature with forward slash which used different isoform letters [3, 2]. For example, HIST1H2BB was known as H2A/f in the earlier nomenclature. The encoded proteins described below are products of genes and transcripts listed in table Table S2. Human canonical core histone polypeptides have typically been numbered with omission of the N-terminal methionine since this is likely to be removed because most sequences have a small hydrophilic amino acid as the second residue [82]. This convention predates the Human Genome Variation Society (HGVS) recommendation to include the initial methionine as residue 1. We have omitted the N-terminal methionine on figures, alignments, and amino-acid numbering for consistency.

### Canonical H2A isoforms

Canonical H2A genes encode 11 different protein isoforms with pairwise identity down to 88% (Table 4). These are separable by TAU PAGE into two bands identified as H2A.1 and H2A.2. TAU PAGE distinguishes leucine from methionine at residue 51 [20, 87], implying there are up to 9 different protein sequences in H2A.1 and 2 in H2A.2. There is no concordance between the bands H2A.1 and H2A.2 and the location of isoform-encoding genes in HIST1 and HIST2 histone gene clusters. For example, the HIST2 cluster contains genes encoding isoforms with both Leu51 and Met51 while HIST1, HIST2, and HIST3 clusters all contain genes encoding H2A isoforms with Leu51. No functional distinction between H2A.1 and H2A.2 has been reported.

**Table 4:**
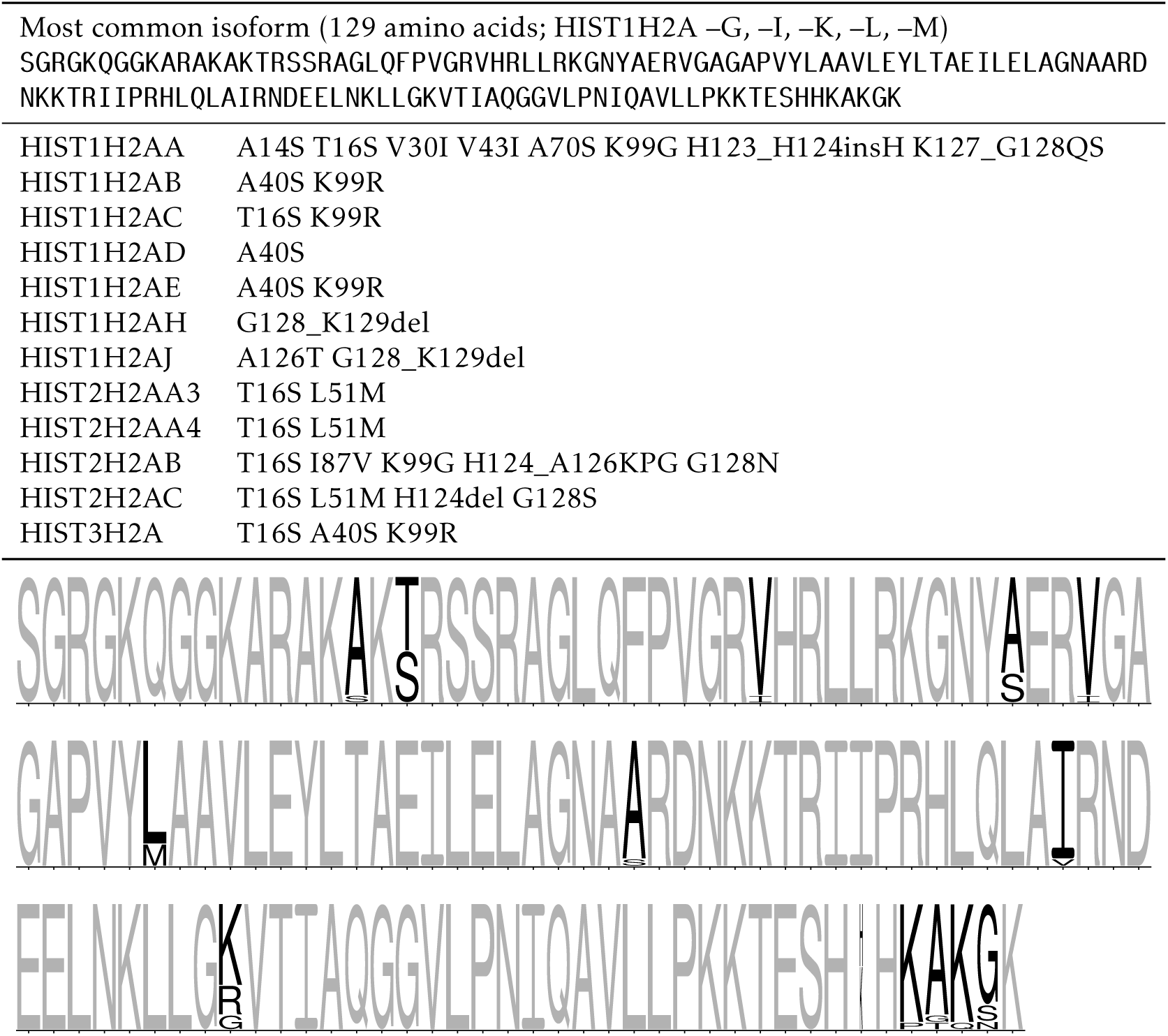
Canonical H2A encoded protein isoforms. Upper panel shows isoform variations relative to the most numerous protein isoform using HGVS recommended nomenclature [13]. Lower panel shows sequence logo of all isoforms aligned with invariant residues in grey.

Excluding *HIST1H2AA* discussed above, the sites of difference in two or more isoforms of canonical H2A are serine or threonine at residue 16, alanine or serine at residue 40, lysine, arginine, or glycine at residue 99, and the C-terminal residues from 124 onwards (Table 4). All these sites have implications for post-translational modifications.

### Canonical H2B isoforms

Canonical H2B has more isoforms than the other histone types (Table 5) with 15 unique proteins diverging down to 77% pairwise identity. Nevertheless, all isoforms appear to migrate together in TAU PAGE.

**Table 5:**
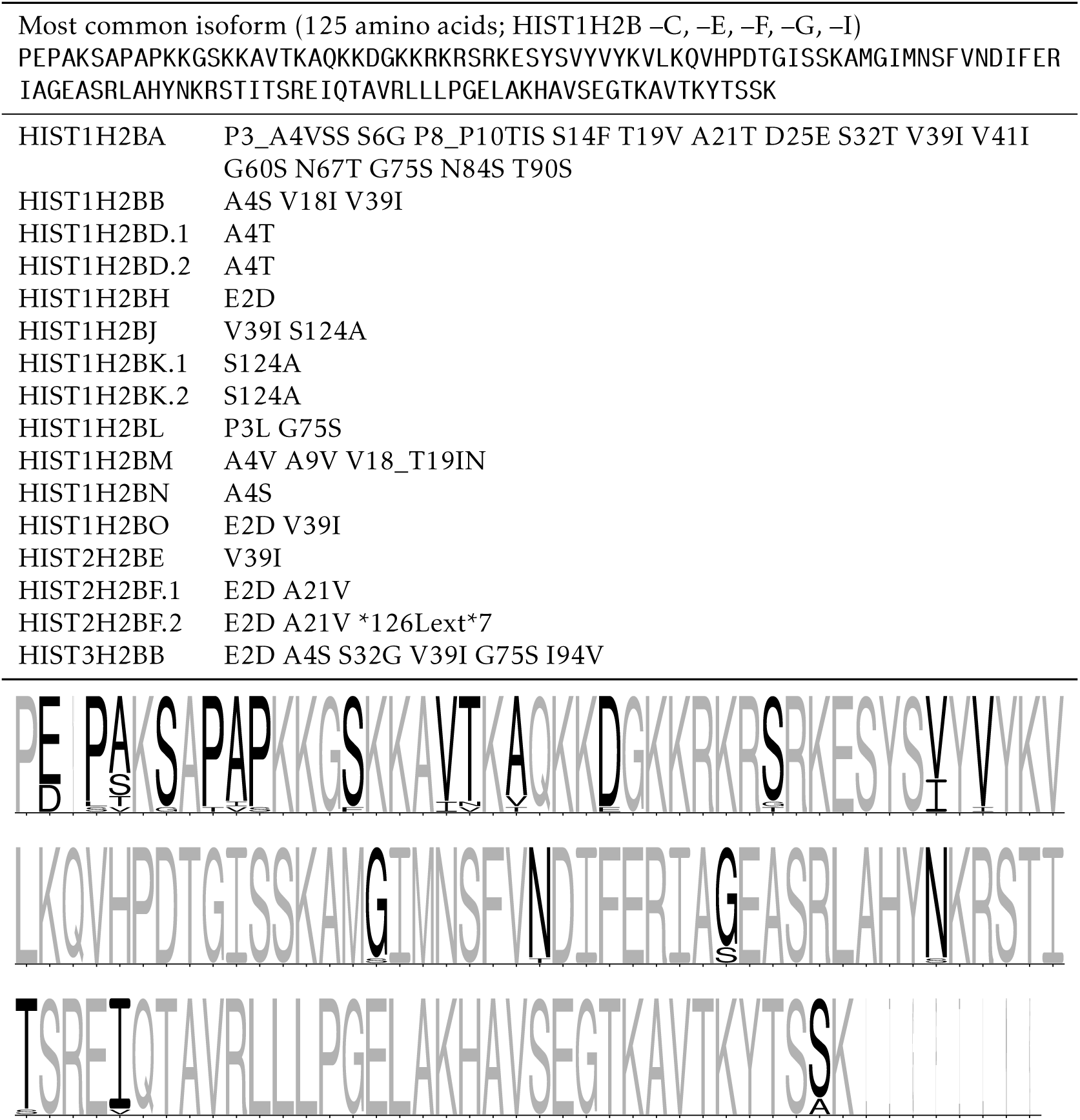
Canonical H2B encoded protein isoforms. Upper panel shows isoform variations relative to the most numerous protein isoform using HGVS recommended nomenclature [14]. For clarity, isoforms encoded by multiple transcripts of a single gene are distingushed by a numerical suffix (see Table S2). Lower panel shows sequence logo of all isoforms aligned with invariant residues in grey.

There is significant variability between isoforms in the N-terminal region and this is one of the most variable sites between canonical core histones of different species. Human H2B has a unique and distinctive N-terminal proline-acidic-proline motif (PEP/PDP) followed by a very variable residue that can be alanine, serine, threonine, or valine (Table 5).

Excluding *HIST1H2BA* discussed above, the remaining isoform differences in two or more isoform are mainly the chemically similar valine or isoleucine at residue 39, and serine to glycine and alanine at residues 75 and 124 respectively, which have have posttranslational modification implications. A number of H2B genes are annotated to have multiple transcripts, although in most cases these transcripts encode identical protein isoforms. Variation in H2B transcript isoform levels between multiple cancer cell lines has been observed [52], although the functional implications are unknown.

### Canonical H3 isoforms

Canonical H3 genes encode only 4 different protein isoforms (Table 6). The majority of H3 genes are in the HIST1 cluster and encode a single polypeptide sequence [16] whereas the canonical H3 genes in HIST2 encode a distinct isoform with the interesting difference of serine instead of cysteine at residue 96 that is separable by TAU PAGE [20]. By apparent coincidence this means HIST1-encoded canonical H3 isoforms are identified as H3.1 and the HIST2-encoded copies are H3.2. As discussed above, *HIST3H3* with four amino acid differences appears to be the largely testes specific variant H3T that has been assigned as H3.4 in variant nomenclature [75] even though it would be predicted to migrate with H3.1 in TAU PAGE [20].

**Table 6:**
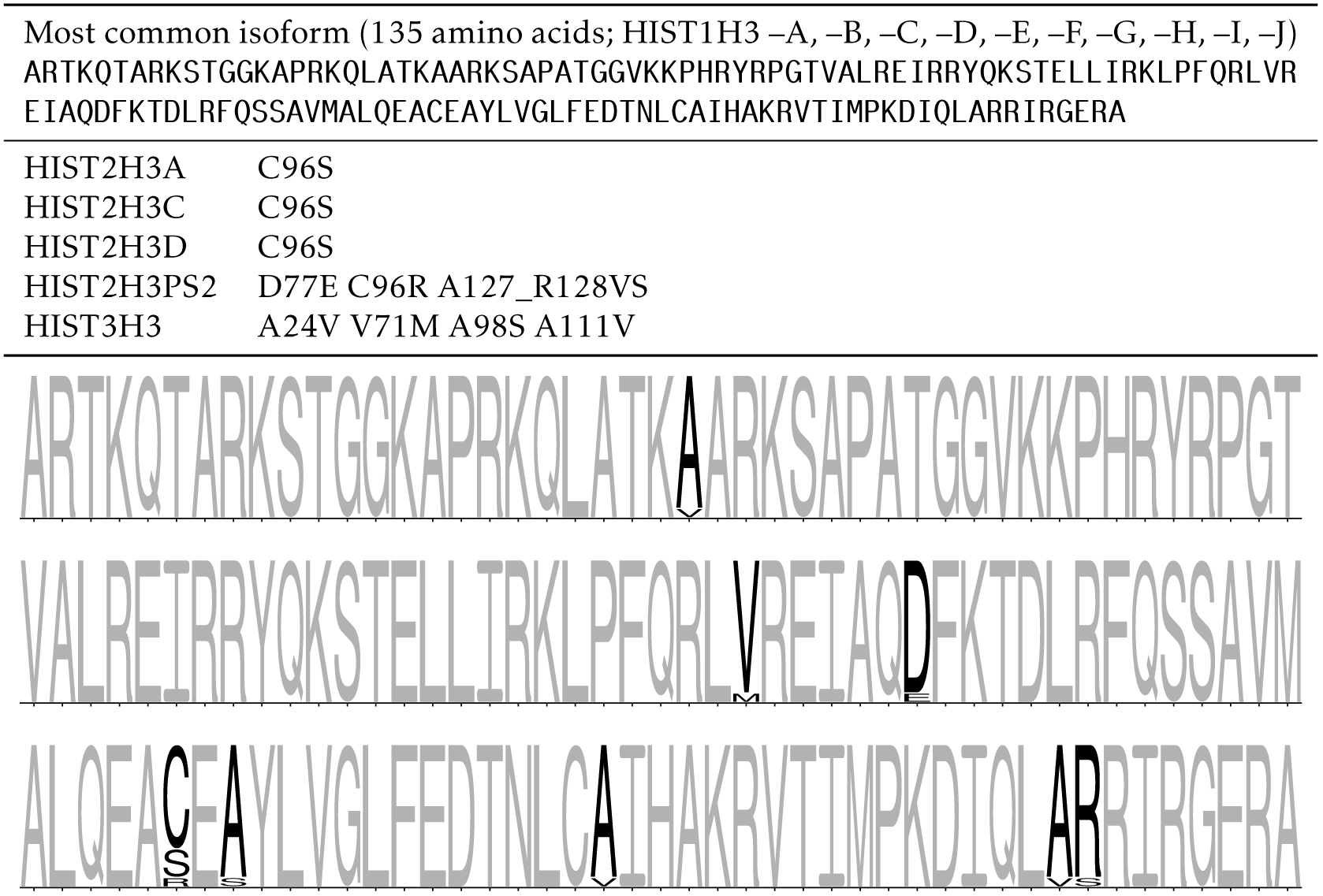
Canonical H3 encoded protein isoforms. Upper panel shows isoform variations relative to the most numerous protein isoform using HGVS recommended nomenclature [13]. Lower panel shows sequence logo of all isoforms aligned with invariant residues in grey.

### Canonical H4 isoforms

H4 is the most homogeneous of all canonical core histones, with all but one of the 15 genes encoding an identical protein sequence (Table 7). These genes are located across HIST1, HIST2, and the isolated *HIST4H4*.

**Table 7:**
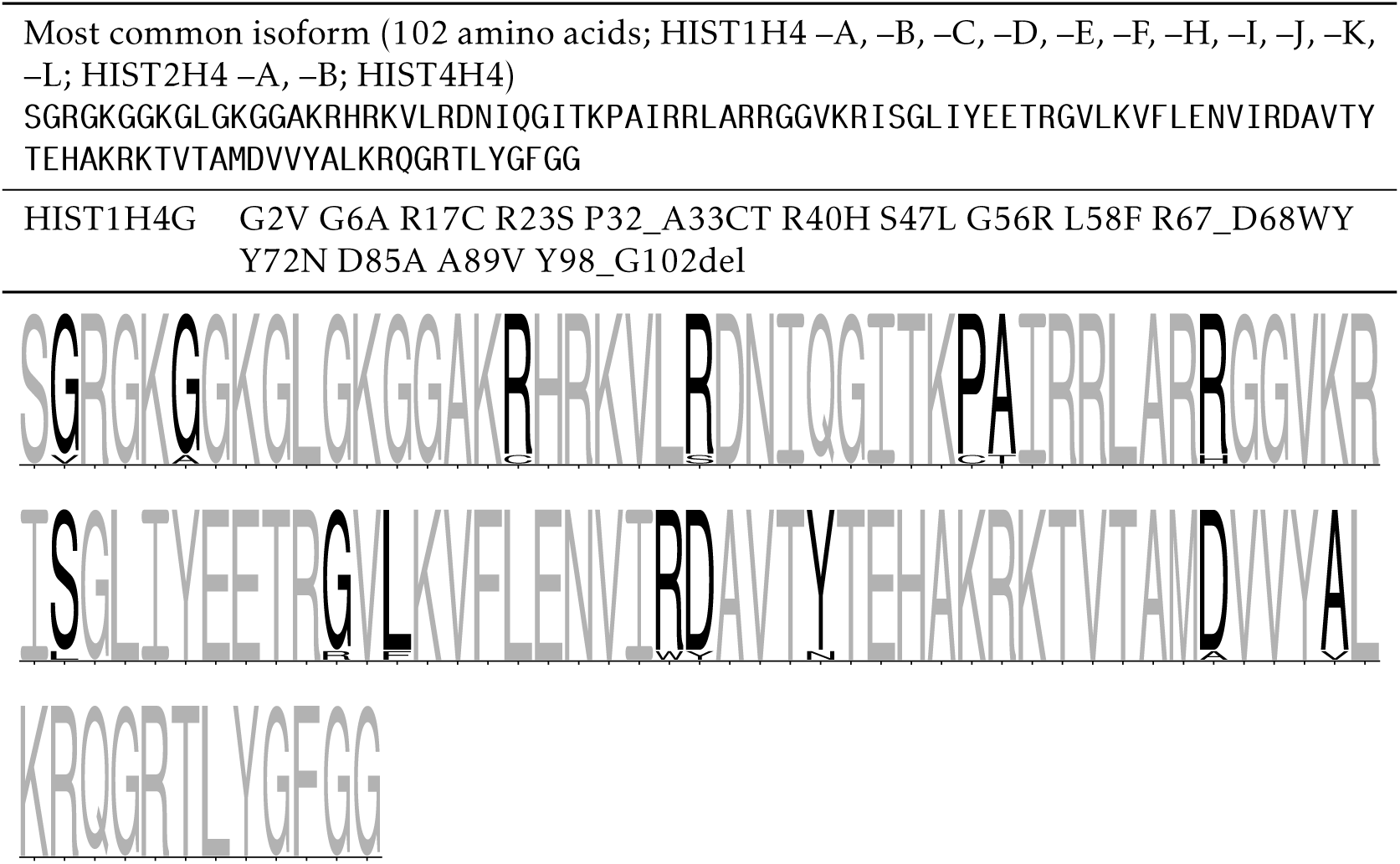
Canonical H4 encoded protein isoforms. Upper panel shows isoform variations relative to the most numerous protein isoform using HGVS recommended nomenclature [13]. Lower panel shows sequence logo of all isoforms aligned with invariant residues in grey.

The divergent *HIST1H4G* gene in the middle of the HIST1 cluster encodes an isoform with 15 amino acid differences and a deletion of the C-terminal 5 residues. This gene is annotated as being transcribed and merits further investigation.

## Reproducible research

A considerable number of builds and updates of the human genome sequence have been made since the last major survey of human canonical histone genes in 2002 [49]. This has resulted in 20 canonical core histone genes added, 3 removed, and 15 sequences being updated for the current RefSeq release (Table S1). These changes reflect the ongoing nature and challenges of genome curation and annotation in reference databases [8].

It is inevitable that sequences and annotations will continue to be revised based on continuous curation and improved experimental insights. This in turn prompts reevaluation of assumptions about their biological contributions, illustrated by the recognition that some genes annotated as canonical core histone isoforms are testes-specific variants [75], and by new observations of cell type-specific expression [52].

It is important for communities of researchers to contribute to ongoing formal curation of genomics resources by feeding back information to database maintainers [70], although many are unaware of this opportunity [33]. In the course of this work we have suggested a significant number of proposals for improvements to RefSeq curators. These were identified automatically by scripted tests for consistency and uniformity between the database annotations and expected properties of histones. Table S11 lists the current apparent anomalies from these tests and is the basis for ongoing discussions with curators.

Dynamic data can be presented in specialist online database resources such as the Histone Database [15] and HIstome [37]. These database interfaces provide comprehensive access to data but have limited curation and are difficult to cite. They can also cease to be updated or become inaccessible. In contrast, manuscripts are the established method of scientific communication because they provide descriptive context and accessible formatting for readers, can be peer reviewed, and have well-established methods for citation. There are established mechanisms for permanent archival of published manuscripts. However, static catalogues in manuscripts such as earlier surveys of histone genes [2, 49] inevitably become supersed by improvements in source data and cannot be updated to remove errors.

A self-updating manuscript bridges the features of dynamic database and static manuscript presentation styles by providing convenient access to the most current information. It encourages communities of researchers to directly feed into the formal curation process and enables rapid leveraging of the most current data for relatively stable gene families.

Implementating a dynamic manuscript is also an example of “reproducible research” [22, 84] that can address the topical challenge of irreproducibility in biological data and interpretation [53, 35].

This manuscript is generated directly from sequences and annotations in the core NCBI RefSeq resource [56]. The processing system for the manuscript does not cache intermediate information, so all changes contributed by the community of histone researchers and curated by professional database maintainers are directly and automatically reflected at each manuscript refresh. All scripts including instructions for automatic builds are transparently available in a public repository. Core processing is based on a BioPerl program contributed to the Bio-EUtilities distribution and publicly available since 2013. Dependencies on raw data sources, alignment algorithms, or display output libraries can also be upgraded since the processing is automatic and script-based.

One potential challenge for researchers is the referencing of dynamic data within such a manuscript. This can be simply addressed by users citing the publication as a traditional static manuscript then stating the build date in Materials and Methods, as they would for a database.

Although the dynamic data will remain current, major revisions in understanding of a gene family or accumulation of small insights can in time render the explanatory static manuscript text obsolete. In this case the manuscript can simply be refreshed and republished independently in the same way as a traditional manuscript would be. The underlying scripts generating the dynamic data do not need to be rewritten, although new functions can be added to reflect new insights.

The core scripts underlying this manuscript have been written to facilitate generating equivalent catalogues for other organisms via simple build options. We have successfully trialled this for *Mus musculus* (not shown) to demonstrate that an equivalent dynamic manuscript for other histone gene sets only requires drafting appropriate surrounding static explanatory text. The approach is also applicable to larger and more diverse gene families such as our previous cataloguing of the Snf2 family [19].

## Conclusion

Comprehensive tabulation and analysis of the canonical core histone gene family reveals significant diversity of isoforms, and challenges assumptions of bulk chromatin homogeneity. It has led to improvements in consistency of genome annotations and provides a reference for interpreting genomic and proteomic datasets. The differences between canonical core histone protein isoforms uncovers novel questions about their functional significance that suggest future experimental investigations. As a dynamic manuscript, this catalogue can automatically generate an up-to-date overview of the human canonical core histone gene family. It also demonstrates the potential for integrating reproducible research approaches into the scientific literature.

## Materials and methods

The primary manuscript is generated from LAT_E_X sources, derived from dynamic data, and built using SCons [38]. To search the Entrez Gene database and download the associated genomic, transcript, and protein sequences, we have created the bp_genbank_ref_extractor program. This program has been included in the Bio-EUtilities distribution since version 1.73. Analysis of the data relies heavily on BioPerl [69].

All sources are freely available in a git repository at https://github.com/af-lab/histone-catalogue.git, including all sources for figures, manuscript templates, and build system making public all parameters used for processing.

This build of the manuscript was generated using BioPerl 1.007002 and Bio-EUtilities 1.75. Sequence and annotation data was obtained from NCBI RefSeq [56] on 25th July 2019. Sequence alignments were generated by T-Coffee 12.00.7 [55]. Sequence logos were generated using WebLogo version 3.6.0 [11]. Description of sequence variants are represented following HGVS recommended nomenclature [14].

## Supporting information

sequence files used in the analysis

## Acknowledgements

We are grateful to the Perl community on the **irc**.**freenode**.**net** channels #perl and #bioperl, especially Altreus, mst, LeoNerd, and pyrimidine. Invaluable support was also provided by the T_E_X stackexchange community, in particular David Carlisle. DMSP acknowledges the support of the Portuguese Foundation for Science and Technology (FCT).

## Supplementary Data

**Supplementary Table S1:**
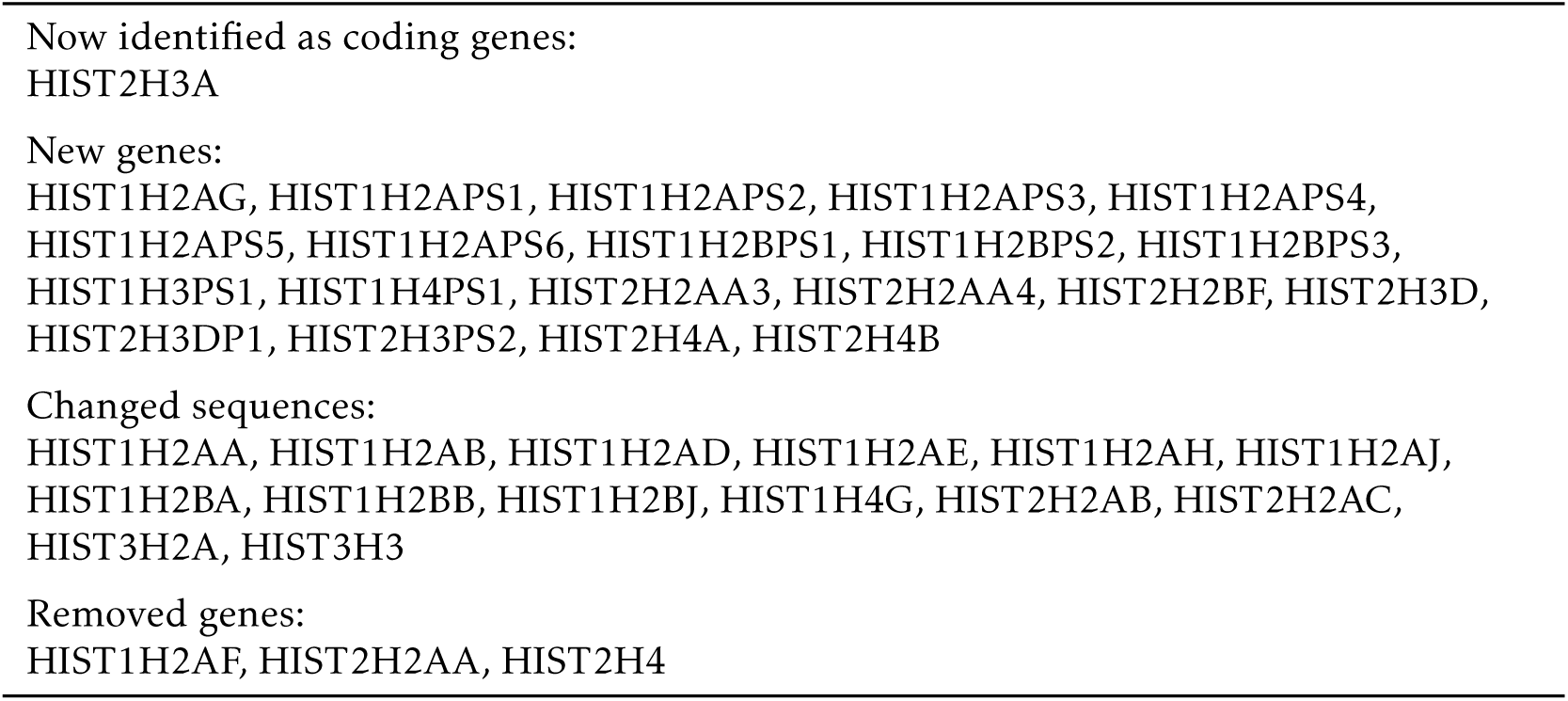
Changes in human canonical core histone gene catalogue annotated in NCBI RefSeq obtained on 25th July 2019 compared to *Marzluff et al*. [49].

**Supplementary Table S2:** Catalogue of annotated human canonical core histone genes, transcripts, and encoded proteins. Data obtained from NCBI RefSeq [56] on 25th July 2019.

| Type | Gene name | Gene UID | Transcript accession | Protein accession |
| --- | --- | --- | --- | --- |
| H2A | HIST1H2AA | 221613 | NM_170745 | NP_734466 |
| H2A | HIST1H2AB | 8335 | NM_003513 | NP_003504 |
| H2A | HIST1H2AC | 8334 | NM_003512 | NP_003503 |
| H2A | HIST1H2AD | 3013 | NM_021065 | NP_066409 |
| H2A | HIST1H2AE | 3012 | NM_021052 | NP_066390 |
| H2A | HIST1H2AG | 8969 | NM_021064 | NP_066408 |
| H2A | HIST1H2AH | 85235 | NM_080596 | NP_542163 |
| H2A | HIST1H2AI | 8329 | NM_003509 | NP_003500 |
| H2A | HIST1H2AJ | 8331 | NM_021066 | NP_066544 |
| H2A | HIST1H2AK | 8330 | NM_003510 | NP_003501 |
| H2A | HIST1H2AL | 8332 | NM_003511 | NP_003502 |
| H2A | HIST1H2AM | 8336 | NM_003514 | NP_003505 |
| H2A | HIST1H2APS1 | 387319 | pseudogene | pseudogene |
| H2A | HIST1H2APS2 | 85303 | pseudogene | pseudogene |
| H2A | HIST1H2APS3 | 387323 | pseudogene | pseudogene |
| H2A | HIST1H2APS4 | 8333 | pseudogene | pseudogene |
| H2A | HIST1H2APS5 | 10341 | pseudogene | pseudogene |
| H2A | HIST1H2APS6 | 100509927 | pseudogene | pseudogene |
| H2A | HIST2H2AA3 | 8337 | NM_003516 | NP_003507 |
| H2A | HIST2H2AA4 | 723790 | NM_001040874 | NP_001035807 |
| H2A | HIST2H2AB | 317772 | NM_175065 | NP_778235 |
| H2A | HIST2H2AC | 8338 | NM_003517 | NP_003508 |
| H2A | HIST3H2A | 92815 | NM_033445 | NP_254280 |
| H2B | HIST1H2BA | 255626 | NM_170610 | NP_733759 |
| H2B | HIST1H2BB | 3018 | NM_021062 | NP_066406 |
| H2B | HIST1H2BC | 8347 | NM_003526 | NP_003517 |
| H2B | HIST1H2BD | 3017 | NM_021063 | NP_066407 |
| H2B | HIST1H2BD | 3017 | NM_138720 | NP_619790 |
| H2B | HIST1H2BE | 8344 | NM_003523 | NP_003514 |
| H2B | HIST1H2BF | 8343 | NM_003522 | NP_003513 |
| H2B | HIST1H2BG | 8339 | NM_003518 | NP_003509 |
| H2B | HIST1H2BH | 8345 | NM_003524 | NP_003515 |
| H2B | HIST1H2BI | 8346 | NM_003525 | NP_003516 |
| H2B | HIST1H2BJ | 8970 | NM_021058 | NP_066402 |
| H2B | HIST1H2BK | 85236 | NM_001312653 | NP_001299582 |
| H2B | HIST1H2BK | 85236 | NM_080593 | NP_542160 |
| H2B | HIST1H2BL | 8340 | NM_003519 | NP_003510 |
| H2B | HIST1H2BM | 8342 | NM_003521 | NP_003512 |
| H2B | HIST1H2BN | 8341 | NM_003520 | NP_003511 |
| H2B | HIST1H2BO | 8348 | NM_003527 | NP_003518 |
| H2B | HIST1H2BPS1 | 100288742 | pseudogene | pseudogene |
| H2B | HIST1H2BPS2 | 10340 | pseudogene | pseudogene |
| H2B | HIST1H2BPS3 | 100820735 | pseudogene | pseudogene |
| H2B | HIST2H2BA | 337875 | pseudogene | pseudogene |
| H2B | HIST2H2BB | 338391 | pseudogene | pseudogene |
| H2B | HIST2H2BC | 337873 | pseudogene | pseudogene |
| H2B | HIST2H2BD | 337874 | pseudogene | pseudogene |
| H2B | HIST2H2BE | 8349 | NM_003528 | NP_003519 |
| H2B | HIST2H2BF | 440689 | NM_001024599 | NP_001019770 |
| H2B | HIST2H2BF | 440689 | NM_001161334 | NP_001154806 |
| H2B | HIST3H2BA | 337872 | pseudogene | pseudogene |
| H2B | HIST3H2BB | 128312 | NM_175055 | NP_778225 |
| H3 | HIST1H3A | 8350 | NM_003529 | NP_003520 |
| H3 | HIST1H3B | 8358 | NM_003537 | NP_003528 |
| H3 | HIST1H3C | 8352 | NM_003531 | NP_003522 |
| H3 | HIST1H3D | 8351 | NM_003530 | NP_003521 |
| H3 | HIST1H3E | 8353 | NM_003532 | NP_003523 |
| H3 | HIST1H3F | 8968 | NM_021018 | NP_066298 |
| H3 | HIST1H3G | 8355 | NM_003534 | NP_003525 |
| H3 | HIST1H3H | 8357 | NM_003536 | NP_003527 |
| H3 | HIST1H3I | 8354 | NM_003533 | NP_003524 |
| H3 | HIST1H3J | 8356 | NM_003535 | NP_003526 |
| H3 | HIST1H3PS1 | 100289545 | pseudogene | pseudogene |
| H3 | HIST2H3A | 333932 | NM_001005464 | NP_001005464 |
| H3 | HIST2H3C | 126961 | NM_021059 | NP_066403 |
| H3 | HIST2H3D | 653604 | NM_001123375 | NP_001116847 |
| H3 | HIST2H3DP1 | 106479023 | pseudogene | pseudogene |
| H3 | HIST2H3PS2 | 440686 | NM_001355409 | NP_001342338 |
| H3 | HIST3H3 | 8290 | NM_003493 | NP_003484 |
| H4 | HIST1H4A | 8359 | NM_003538 | NP_003529 |
| H4 | HIST1H4B | 8366 | NM_003544 | NP_003535 |
| H4 | HIST1H4C | 8364 | NM_003542 | NP_003533 |
| H4 | HIST1H4D | 8360 | NM_003539 | NP_003530 |
| H4 | HIST1H4E | 8367 | NM_003545 | NP_003536 |
| H4 | HIST1H4F | 8361 | NM_003540 | NP_003531 |
| H4 | HIST1H4G | 8369 | NM_003547 | NP_003538 |
| H4 | HIST1H4H | 8365 | NM_003543 | NP_003534 |
| H4 | HIST1H4I | 8294 | NM_003495 | NP_003486 |
| H4 | HIST1H4J | 8363 | NM_021968 | NP_068803 |
| H4 | HIST1H4K | 8362 | NM_003541 | NP_003532 |
| H4 | HIST1H4L | 8368 | NM_003546 | NP_003537 |
| H4 | HIST1H4PS1 | 10337 | pseudogene | pseudogene |
| H4 | HIST2H4A | 8370 | NM_003548 | NP_003539 |
| H4 | HIST2H4B | 554313 | NM_001034077 | NP_001029249 |
| H4 | HIST4H4 | 121504 | NM_175054 | NP_778224 |

**Supplementary Table S3:**
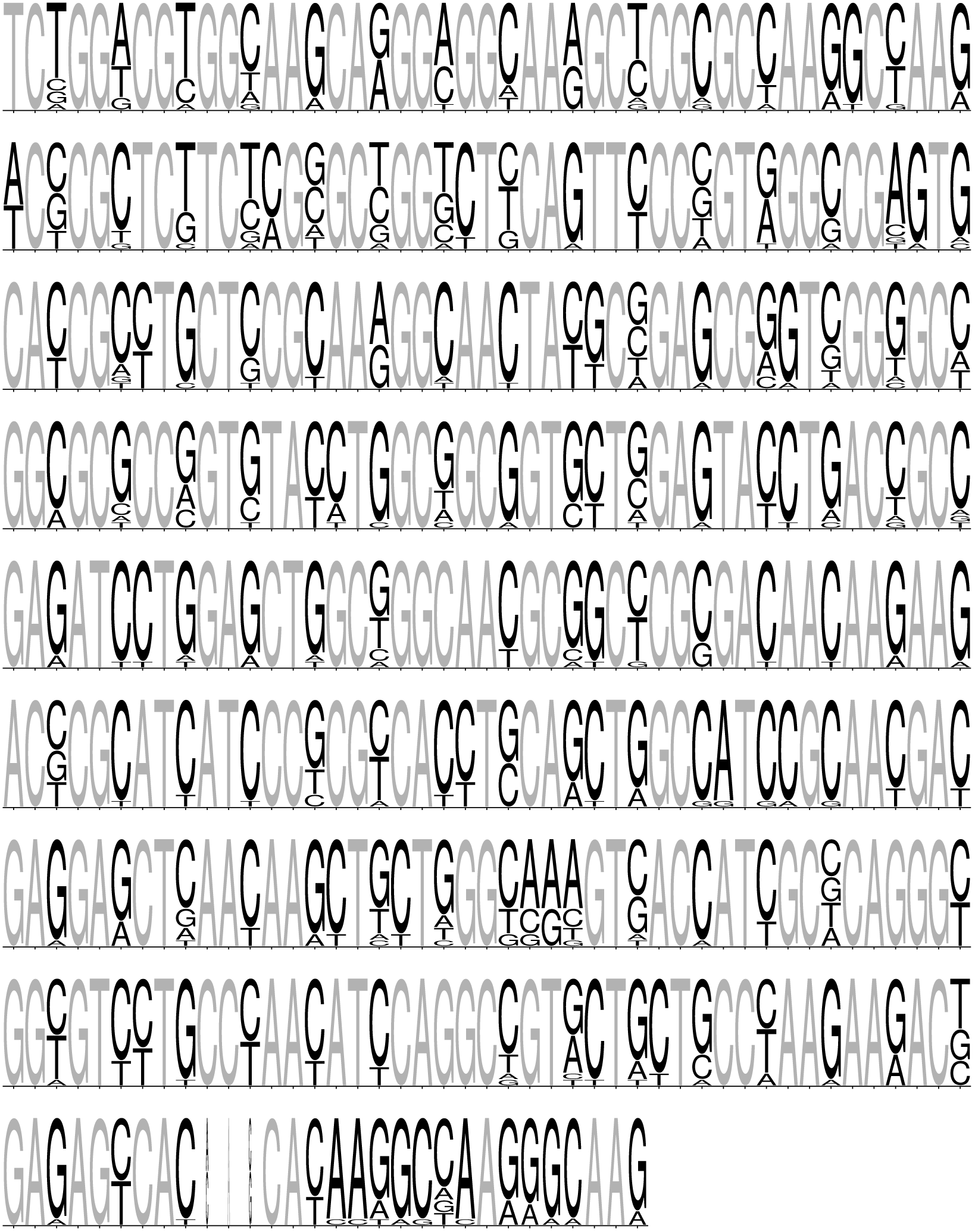
Sequence logo for the human canonical H2A gene coding regions listed in table Table S2. Initiator codon ATG and stop codon are omitted.

**Supplementary Table S4:**
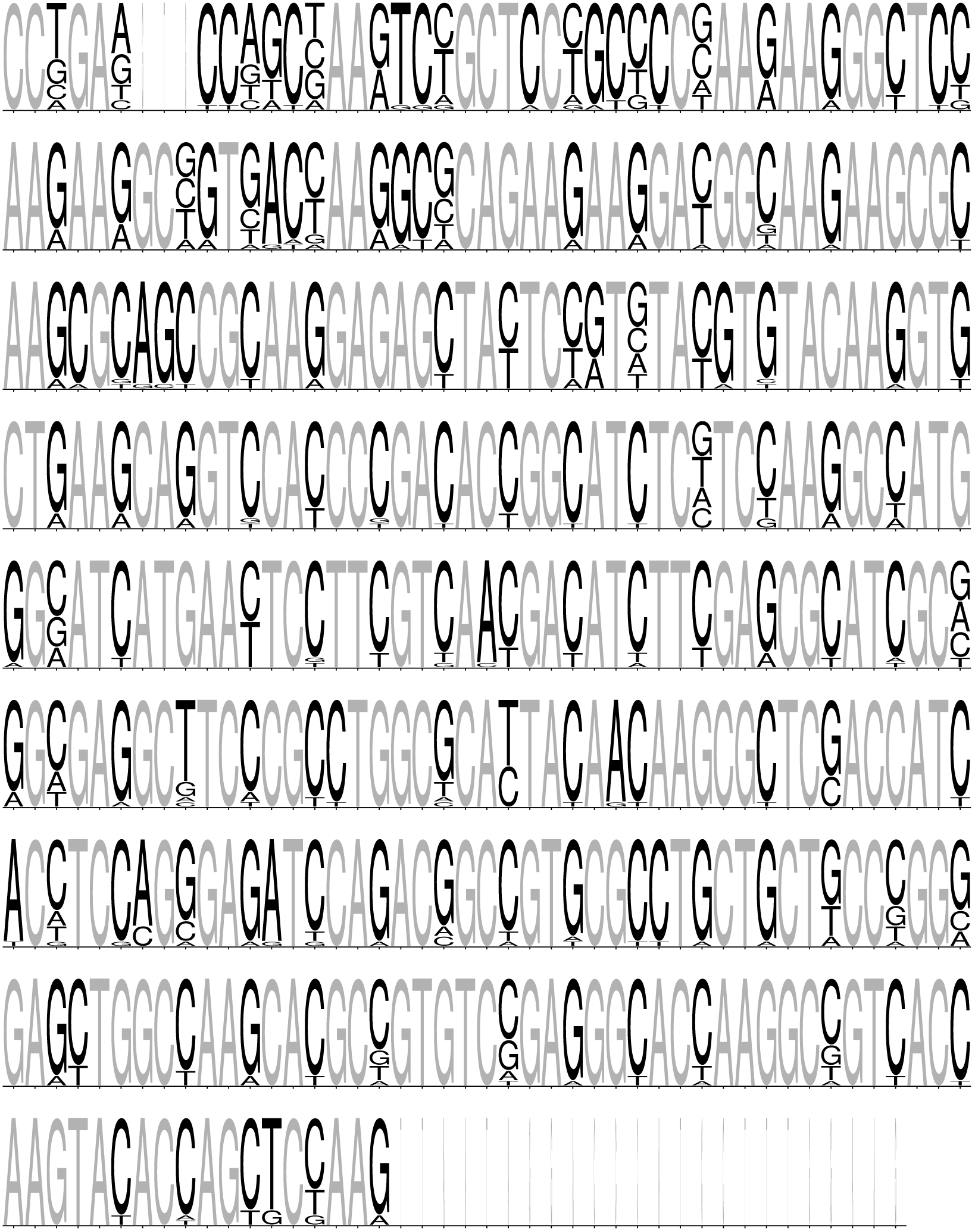
Sequence logo for the human canonical H2B gene coding regions listed in table Table S2. Initiator codon ATG and stop codon are omitted.

**Supplementary Table S5:**
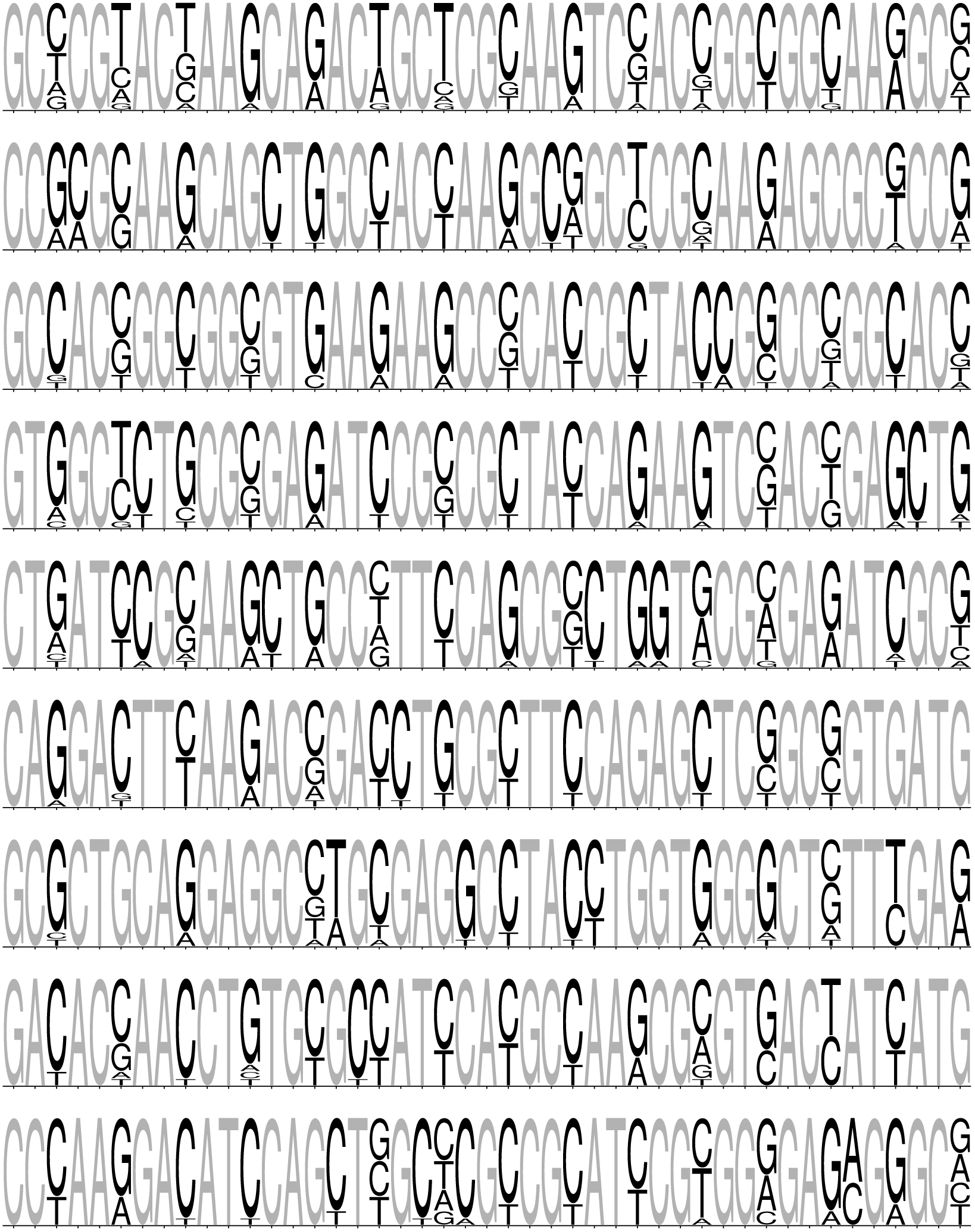
Sequence logo for the human canonical H3 gene coding regions listed in table Table S2. Initiator codon ATG and stop codon are omitted.

**Supplementary Table S6:**
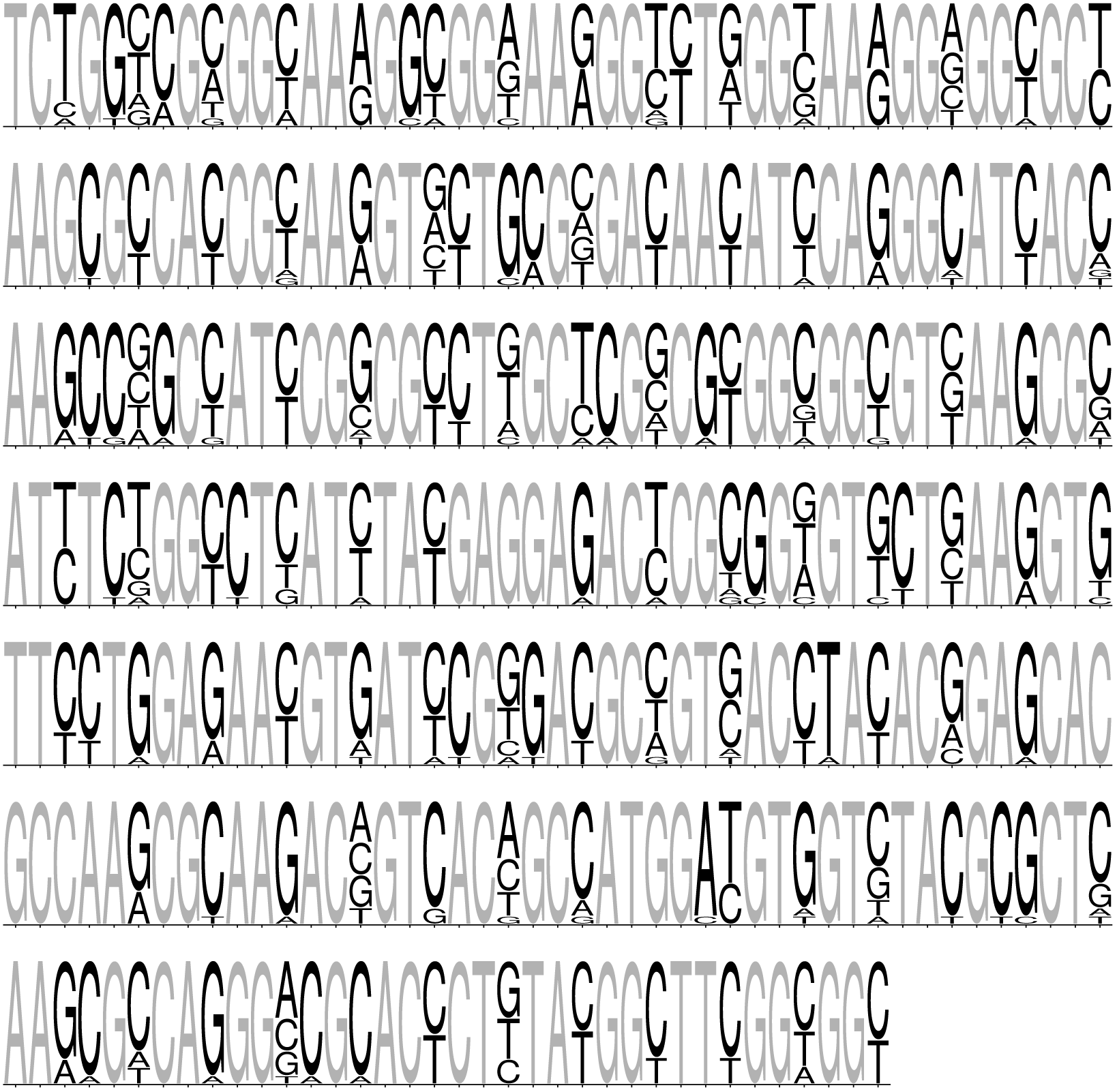
Sequence logo for the human canonical H4 gene coding regions listed in table Table S2. Initiator codon ATG and stop codon are omitted.

**Supplementary Table S7:** Annotated histone variants. HGNC histone family names [27], commonly used protein names, and nomenclature proposal of *Talbert et al*. [75].

| Family | Common name | *Talbert et al.* [75] | Notes |
| --- | --- | --- | --- |
| H2AFB | H2A.Bbd | H2A.B | equivalent to H2A.L |
| H2AFJ | H2A.J | H2A.J | at HIST4 locus |
| H2AFV | H2A.Z-2 | H2A.Z.2 | - |
| H2AFX | H2AX/H2A.X | H2A.X | - |
| H2AFY | macroH2A/mH2A | macroH2A | - |
| H2AFZ | H2A.Z | H2A.Z.1 | - |
| H2BFM | ? | H2B.M | homologous to H2B.W,<br>X-linked |
| H2BFS | - | - | pseudogene |
| H2BFWT | ? | H2B.W | testes specific, X-linked |
| H2BFX | - | - | pseudogene |
| HIST1H2BA | TH2B/TSH2B | TS H2B.1 | testes specific |
| HIST3H3 | H3T | H3.4 | testes specific |
| H3F3 | H3.3 | H3.3 | euchromatin related |
| CENPA | CENP-A | - | centromere-specific |

**Supplementary Table S8:** Mean synonymous and non-synonymous distances between pairs of canonical core histone genes. Mean over all pairwise comparisons of non-synonymous (*d*_*N*_) and synonymous (*d*_*S*_) nucleotide substitutions and mean of *d*_*N*_ */d*_*S*_ ratios for canonical core histone coding regions computed using *Goldman and Yang* [24] model implemented in codeml from the PAML package [58].

| | $d_N$ | $d_S$ | $d_N/d_S$ |
| --- | --- | --- | --- |
| H2A | 0.0113 | 3.5345 | 0.00531 |
| H2B | 0.0159 | 1.7303 | 0.00805 |
| H3 | 0.0053 | 2.6275 | 0.0084 |
| H4 | 0.0134 | 2.7566 | 0.00674 |

**Supplementary Table S9:**
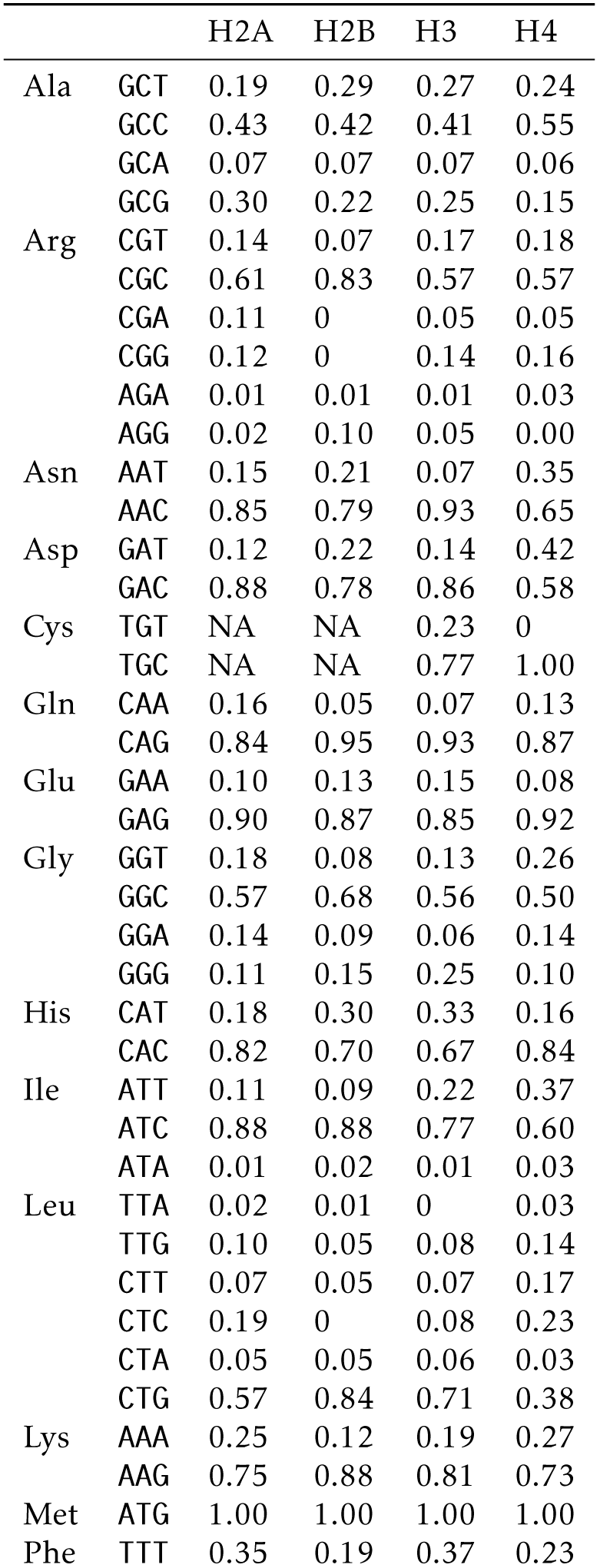
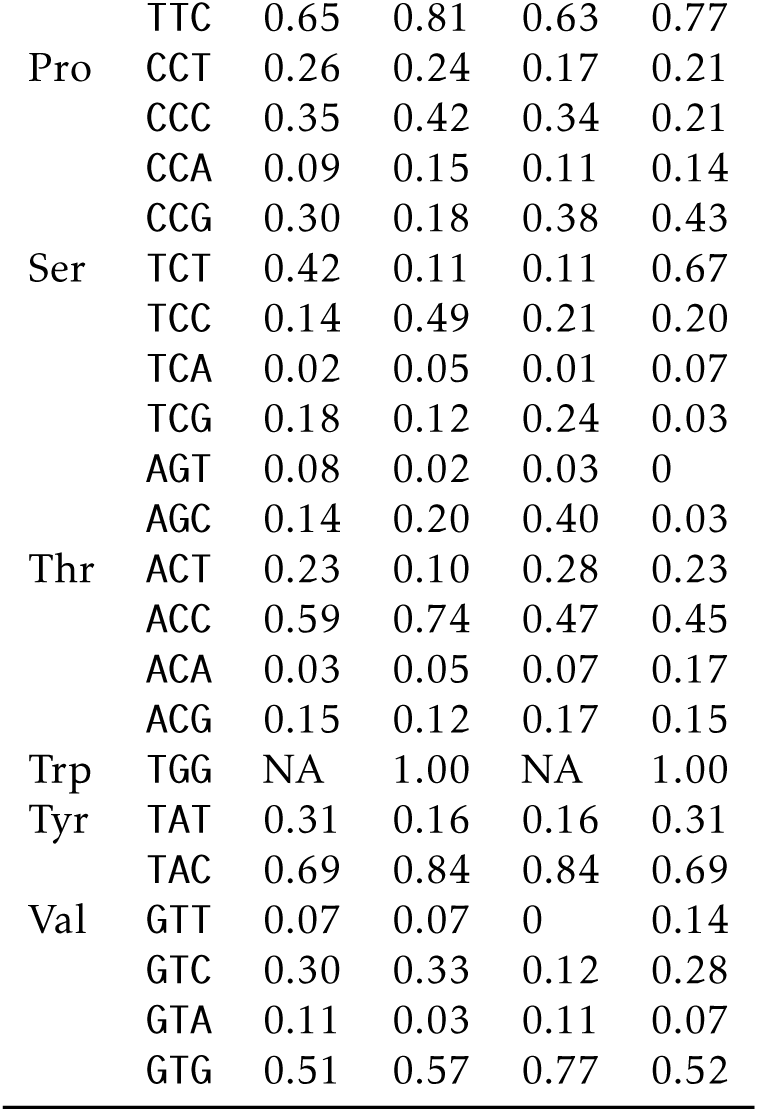
Codon usage frequency for each amino acid and canonical core histone type.

**Supplementary Table S10:**
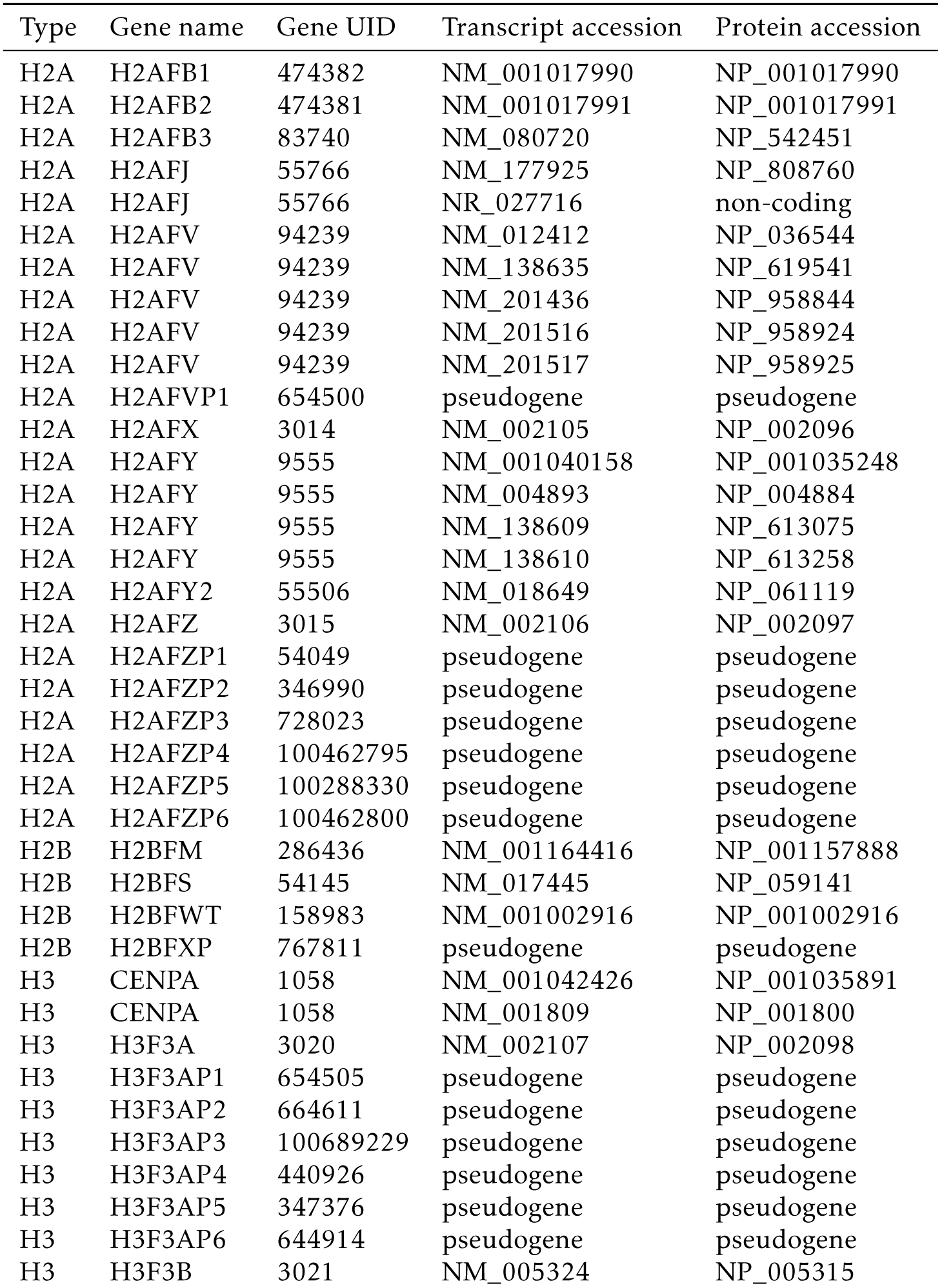
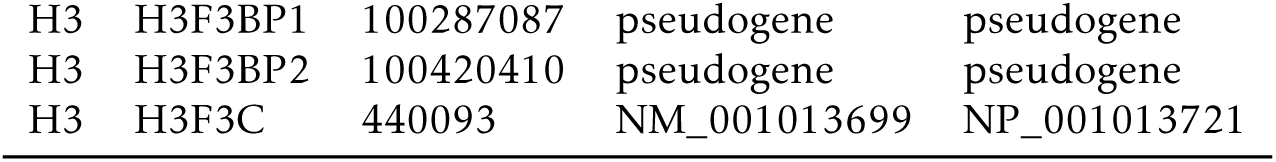
Catalogue of annotated human core histone variant genes, transcripts and encoded proteins. Catalogue of annotated human core histone variant genes, transcripts, and encoded proteins. Data obtained from NCBI RefSeq [56] on 25th July 2019.

| Type | Gene name | Gene UID | Transcript accession | Protein accession |
| --- | --- | --- | --- | --- |
| H2A | H2AFB1 | 474382 | NM_001017990 | NP_001017990 |
| H2A | H2AFB2 | 474381 | NM_001017991 | NP_001017991 |
| H2A | H2AFB3 | 83740 | NM_080720 | NP_542451 |
| H2A | H2AFJ | 55766 | NM_177925 | NP_808760 |
| H2A | H2AFJ | 55766 | NR_027716 | non-coding |
| H2A | H2AFV | 94239 | NM_012412 | NP_036544 |
| H2A | H2AFV | 94239 | NM_138635 | NP_619541 |
| H2A | H2AFV | 94239 | NM_201436 | NP_958844 |
| H2A | H2AFV | 94239 | NM_201516 | NP_958924 |
| H2A | H2AFV | 94239 | NM_201517 | NP_958925 |
| H2A | H2AFVP1 | 654500 | pseudogene | pseudogene |
| H2A | H2AFX | 3014 | NM_002105 | NP_002096 |
| H2A | H2AFY | 9555 | NM_001040158 | NP_001035248 |
| H2A | H2AFY | 9555 | NM_004893 | NP_004884 |
| H2A | H2AFY | 9555 | NM_138609 | NP_613075 |
| H2A | H2AFY | 9555 | NM_138610 | NP_613258 |
| H2A | H2AFY2 | 55506 | NM_018649 | NP_061119 |
| H2A | H2AFZ | 3015 | NM_002106 | NP_002097 |
| H2A | H2AFZP1 | 54049 | pseudogene | pseudogene |
| H2A | H2AFZP2 | 346990 | pseudogene | pseudogene |
| H2A | H2AFZP3 | 728023 | pseudogene | pseudogene |
| H2A | H2AFZP4 | 100462795 | pseudogene | pseudogene |
| H2A | H2AFZP5 | 100288330 | pseudogene | pseudogene |
| H2A | H2AFZP6 | 100462800 | pseudogene | pseudogene |
| H2B | H2BFM | 286436 | NM_001164416 | NP_001157888 |
| H2B | H2BFS | 54145 | NM_017445 | NP_059141 |
| H2B | H2BFWT | 158983 | NM_001002916 | NP_001002916 |
| H2B | H2BFXP | 767811 | pseudogene | pseudogene |
| H3 | CENPA | 1058 | NM_001042426 | NP_001035891 |
| H3 | CENPA | 1058 | NM_001809 | NP_001800 |
| H3 | H3F3A | 3020 | NM_002107 | NP_002098 |
| H3 | H3F3AP1 | 654505 | pseudogene | pseudogene |
| H3 | H3F3AP2 | 664611 | pseudogene | pseudogene |
| H3 | H3F3AP3 | 100689229 | pseudogene | pseudogene |
| H3 | H3F3AP4 | 440926 | pseudogene | pseudogene |
| H3 | H3F3AP5 | 347376 | pseudogene | pseudogene |
| H3 | H3F3AP6 | 644914 | pseudogene | pseudogene |
| H3 | H3F3B | 3021 | NM_005324 | NP_005315 |
| H3 | H3F3BP1 | 100287087 | pseudogene | pseudogene |
| H3 | H3F3BP2 | 100420410 | pseudogene | pseudogene |
| H3 | H3F3C | 440093 | NM_001013699 | NP_001013721 |

**Supplementary Table S11:**
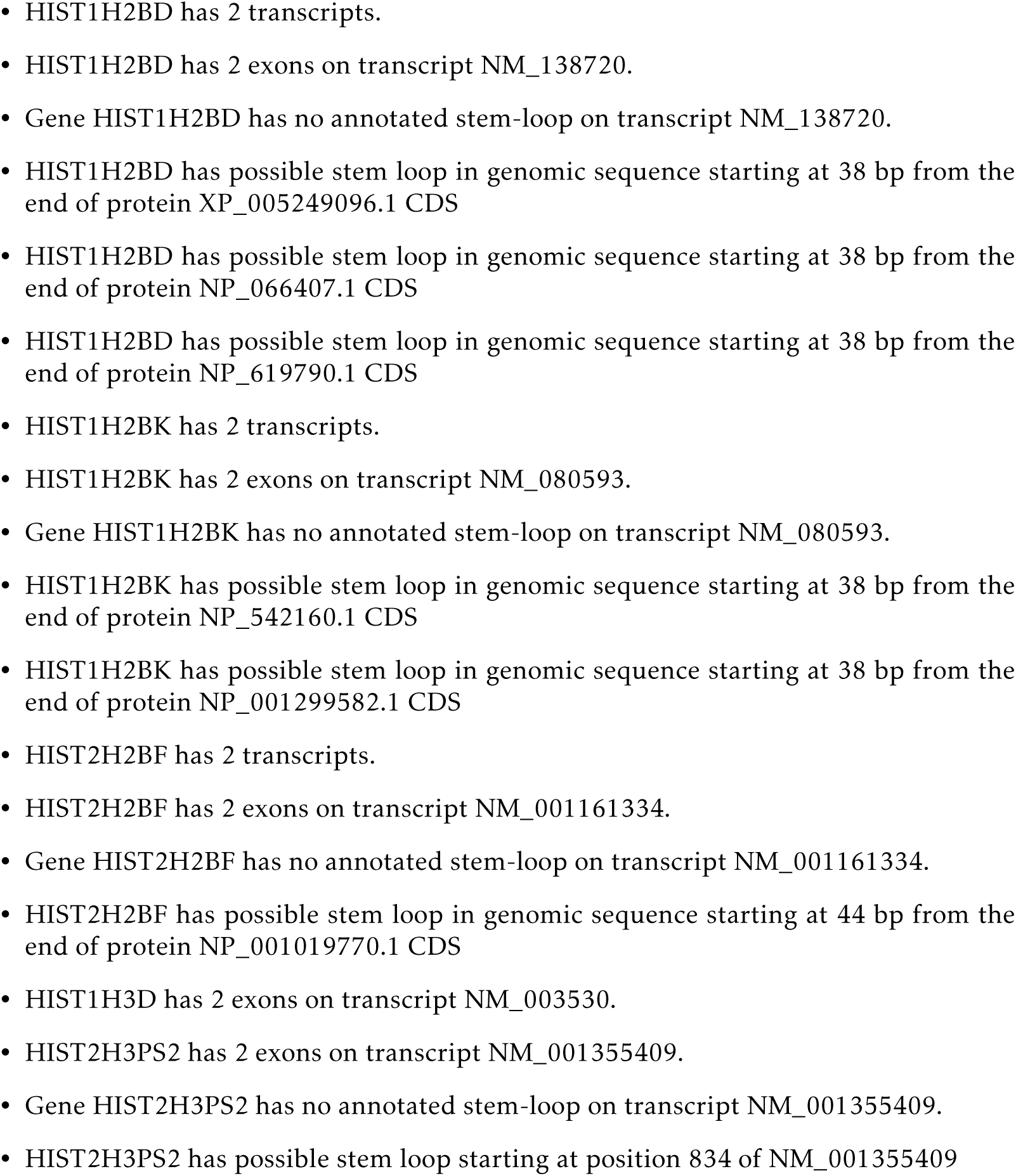
Variations in canonical core histone gene annotations compared to expectation of single exon, single transcript, and stem-loop. Data obtained from NCBI RefSeq [56] on 25th July 2019.

